# Aging, computation, and the evolution of neural regeneration processes

**DOI:** 10.1101/780163

**Authors:** Aina Ollé-Vila, Luís F Seoane, Ricard Solé

## Abstract

Metazoans gather information from their environments and respond in predictable ways. These computational tasks are achieved with neural networks of varying complexity. Their performance must be reliable over an individual’s lifetime while dealing with the shorter lifespan of cells and connection failure – thus rendering aging a relevant feature. How do computations degrade over an organism’s lifespan? How reliable can they remain throughout? We tackle these questions with a multiobjective optimization approach. We demand that digital organisms equipped with neural networks solve a computational task reliably over an extended lifespan. Neural connections are costly (as an associated metabolism in living beings). They also degrade over time, but can be regenerated at some expense. We investigate the simultaneous minimization of both these costs and the computational error. Pareto optimal tradeoffs emerge with designs displaying a broad range of solutions: from small networks with high regeneration rate, to large, redundant circuits that regenerate slowly. The organism’s lifespan and the external damage act as evolutionary pressures. They improve the exploration of the space of solutions and impose tighter optimality constraints. Large damage rates can also constrain the space of possibilities, forcing the commitment of organisms to unique strategies for neural systems maintenance.

## I. INTRODUCTION

A major revolution in the history of multicellular life was the emergence of neurons, a new class of cells that enhanced the processing and storage of information beyond the genetic level [1]. Such revolution enabled fast adaptation to environmental fluctuations. Combining these apt building blocks, a more short-sighted sensor-actuator logic was soon backed by more complex networks, resulting in yet further increased capabilities for information processing and representation of the external environment [2]. Alongside, organism size and longevity also increased. Both required regeneration mechanisms that would sustain organismal coherence over extended periods of time, far beyond the cellular lifespan. An alternative (or complementary) way to guarantee reliable computation is through an appropriate connectivity pattern – e.g. faulty functioning due to loss of connections could be counterbalanced by redundant wires, among other mechanisms. How this can be achieved was early explored by the first generation of mathematicians dealing with unreliable computers [3, 4]. More recent works have addressed some of these questions from the perspective of reliability theory [5], particularly in relation to the role played by parallel versus sequential topologies [6].

The ability to regenerate the nervous system presents an extraordinary variation in Metazoa. Regarding vertebrate species, all of them can produce new neurons postnatally in specific regions of their nervous system, but only some lower vertebrates (fish and amphibians) can significantly repair several neural structures. Some regenerative ability, however, is found also in reptiles and birds, and even in mammals [7]. Remarkably, replacement of all or part of the nervous system has been documented in a few invertebrate phyla, including coelenterates, flatworms, annelids, gastropods and tunicates [8]. Such re-grown neural networks are parsimoniously integrated within the rest of the circuitry, stressing how phenotypic functionality is recovered. Although this ability is largely absent in higher vertebrates, evidence piles up that the potential might lay dormant [9–11]. Another open question concerns whether new neurons are being created throughout our lives in the absence of damage. While few or no new neurons seem to grow in most parts of human brains, there is evidence of limited neurogenesis in the hippocampus [12, 13] and the olfactory bulb [14]. Several species, notably fish, present large rates of neurogenesis, often requiring apoptosis of older cells to make place for the new ones [15].

The origin of this diversity of strategies around nervous system maintenance is a major challenge [16]. What are the evolutionary drivers behind each solution? The metabolic cost of wires readily comes to mind, and has already been explored as a major constraint of neural architecture [17–19], while the regeneration costs are also obvious. The relevance of these factors must be analyzed under the light of a phenotypic function. This, in neural circuits, traces back to computations that must be implemented within reasonable error bounds. This computational performance constitutes a third dimension relevant to our research questions.

What is the optimal tradeoff between these factors for reliable neural circuits? Is the range of solutions parsimonious, or are there more locally stable designs that hinder the access to other possibilities? In order to answer these questions, we asses the maintenance of reliable computations over extended lifespans while enduring an aging process (inflicted through an external damage). Interestingly, the interplay between the lifespan of the whole organism versus the time scale of its constituent parts brings in an extra factor in our study. Its relevance becomes apparent in the empirical record, notably in the apoptosis of older neurons sought by some fish species [15] – implying that a valid regeneration strategy actively shortens the useful life of the organism’s building blocks. How are design spaces affected by these different factors – computational performance, organismal lifespans (and its relation to component time scales), external damages, and the severity of metabolic costs? Early and current research has studied how given computational functions are implemented by evolved neural networks [20–25], but these studies seldom connect wiring costs with possible repair processes.

As noted above, our research questions can only be addressed given a computational task that the underlying circuitry must solve. In living organisms this further results in diverse anatomical patterns and neural plans, with the computational tasks ingrained in each organism’s phenotype. This additional diversity falls beyond the scope of this paper. To gain some insight in the tradeoffs behind neural circuits maintenance, we resort to minimal toy models based on networks of Boolean units that solve archetypal tasks. Simple Boolean models have been used to explore the basic principles of neural functions and the role played by architecture, including information propagation thresholds [26], locomotion and gating [27–29], pattern formation [30], or the emergence of modularity [31]. In a more general context of evolved circuits for artificial agents, evolved neural networks play a central role in the development of biologically inspired robotic systems [32, 33].

In this spirit, we test feed-forward networks of Boolean units. We set a fixed number of layers, a varying number of units in each layer and connections across layers, and a range of regeneration rates. These toy neural circuits are tasked with solving a series of computations while responding to the three evolutionary forces outlined above: a cost stemming from computational errors, ii) another cost associated to wiring, and iii) the cost associated to the regeneration of damaged structures. To integrate these evolutionary forces without introducing unjustified biases that would assign more importance to a factor over the others, we adopt a (Pareto) Multi-Objective Optimization (MOO) approach [34–36]. This framework explores designs that simultaneously minimize all costs involved. The solution of a MOO problem is a restricted region in the space of possible designs. This solution embeds the diversity of somehow optimal strategies in a mathematical object whose geometry has been linked to phase transitions [37–41] and criticality [39, 41]. These phenomena give us some insights about how accessible the range of optimal solutions are: whether evolutionary biases can navigate them smoothly as they vary, or whether locally optimal designs dominate under discrete value regions of the different costs so that changes only happen abruptly.

In section II we introduce the elements of our toy model: i) the computational tasks explored, ii) the implementation of the feed-forward networks and their aging process, and iii) how all of this comes together under MOO. In section III we go over the results, including how each evolutionary pressure affects the shape of MOO solutions in design space, and how this relates to the biology of the problem. Section IV concludes discussing our results within existing literature.

## II. METHODS

### A. Computational tasks

We ask our toy neural networks to compute some arbitrary Boolean function:

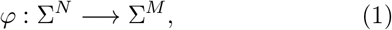

with Σ = {0, 1}. In this paper we used *N* = 3, *M* = 1, and two archetypal computations (figure 1): i) The *multiplexer* (*Mux*) where the “selector” bit *i*_3_ chooses one of the other inputs (*i*_1_ or *i*_2_) as output. ii) The *majority rule* (*Maj*), which returns 1 if most input bits are 1 and 0 otherwise. *Maj* is well known within the study of circuit redundancy and error correction.

**FIG. 1:**
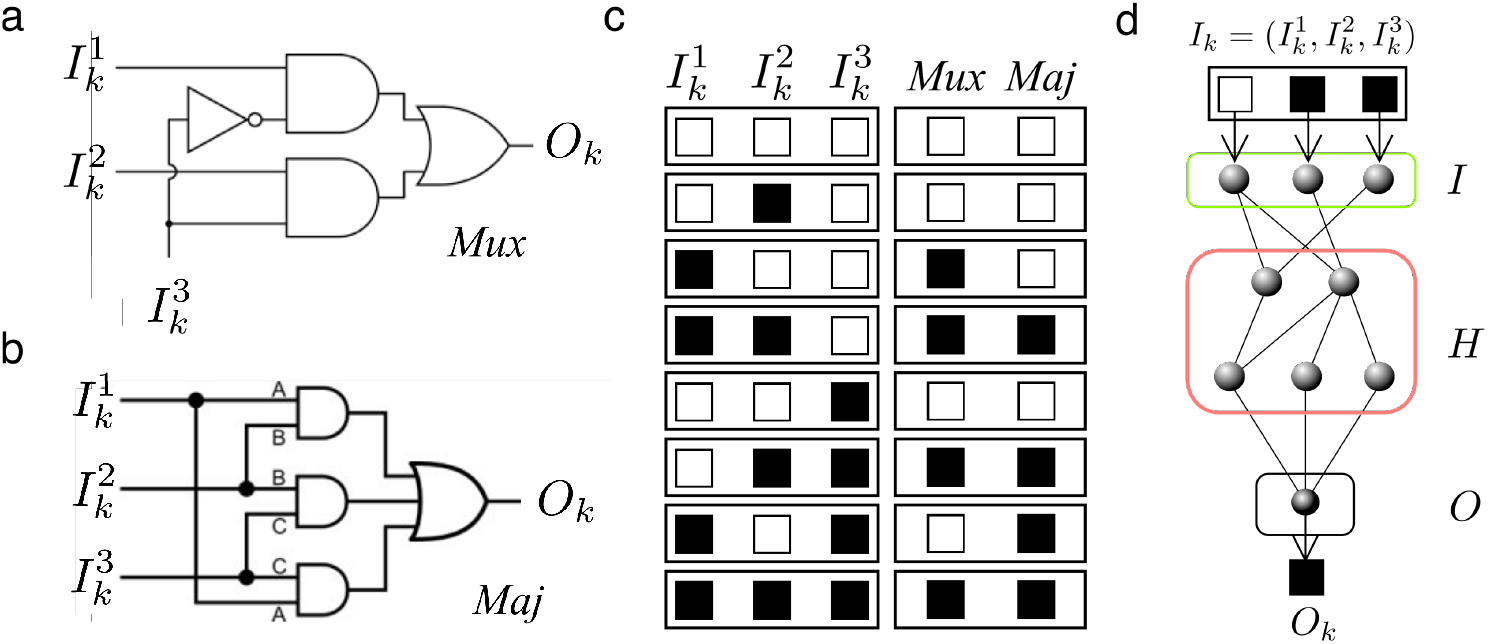
Computational tasks and neural implementation. The agents used in our evolutionary algorithm are tested with two different, three-input, one output logic functions: (a) the multiplexer (*Mux*) and (b) the majority rule (*Maj*). These are traditionally implemented using logic gates as shown. Each circuit has an associated logic table that fully defines each performed computation. (c) Left, the eight possible binary sets of inputs ■ = 0 and □ = 1. Each possible input string, such as ■■□, results in a unique output. The outputs for *Mux* and *Maj* are shown (right). (d) In our evolving system, Boolean gates are replaced by feed-forward neural networks with several layers including input (I), output (O), and hidden (H) ones.

### B. Feed-forward neural networks

We subjected a population of Boolean, feed-forward neural networks to an evolutionary process that optimizes diverse design aspects according to the MOO logic detailed below. Here we clarify the general network architecture, how they compute, and some of the diversity allowed.

Each network consisted of three input units (*I* ≡ {*i*_1_, *i*_2_, *i*_3_}), two hidden layers with a varying number of units in each layer (from 1 to *h_max_* = 15) and connections across layers, and one output unit (*O* ≡ {*o*_1_}). Each connection carried a weight *ω_ij_* = ±1, and each unit had an activation threshold *θ_i_* ∈ [0.01, 1]. Networks computed following a McCulloch-Pitts function [42]:

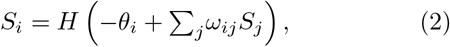

where *j* runs over input signals to unit *i*, and *H*(·) represents the Heaviside step function.

Thus each network computes a Boolean function. With the variety allowed across networks (different number of units, connections, and *θ_i_*), there are several ways to implement a desired function. Different network designs will incur in different costs due to the expense of wiring or regeneration (see below), and their computational reliability will degrade differently for distinct topologies as they *age* (also, see below). This constitutes the basis of our MOO exploration.

### C. Aging process

The networks are evaluated over an extended period *t* = 1, …, *τ* during which their connections are eroded. *τ* defines the required durability for the whole network, which would correspond to the lifespan of a modeled organism. At the beginning of each evaluation step, each connection is removed with a probability *δ*, defining a *damage rate*. Also, the network restores each missing link with probability *ρ*, thus defining a *regeneration rate*. Recovered connections display their original weight (i.e. some detailed memory is never lost).

After knocking off some connections and regenerating others, we assess the reliability of each network in computing *φ* (eq. 1 i.e. *Mux* or *Maj*). An approximate mean-field model (see Supplementary Material, section II) shows how the damage/regeneration process eventually results in an average steady number of missing connections.

We performed experiments with different *δ* and *τ*. Notice that *δ* defines a damage rate, but also contributes to setting an average lifespan (*τ_link_* ~ 1/*δ*) for the network connections. This, together with the timescale of the network lifespan can be combined into a dimensionless ratio *r*_*τ*_ ≡ *δ* · *τ* ~ *τ*/*τ*_*link*_. This ratio is an informative index in analyzing some results, but note that the actual *τ_link_* is also affected by regeneration values.

This aging process might result in plainly unfeasible networks – i.e. graphs that become disconnected such that information cannot flow from input to output. We track the proportion of time that a network *γ* is thus broken through a *feasibility function F*_*f*_(*γ*) ∈ [0, 1]:

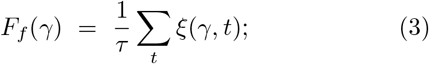

where *ξ*(*γ, t*) = 1 if *γ* remains feasible (connections exist from input to output, and from all input units to the first hidden layer after damage and regeneration at *t*). Otherwise, *ξ*(*γ, t*) = 0 (check section III-E in Supplementary Material for more details).

### D. Multiobjective Optimization

Given the set Γ of all allowed networks (*γ* ∈ Γ), we seek the subset Π ⊂ Γ of designs *γ^π^* ∈ Π that simultaneously minimize all relevant costs without introducing artificial biases. This subset (Π) is the solution of a MOO or Pareto optimization problem [34–36]. Pareto optimal networks *γ^π^* ∈ Π are such that you cannot find two of them 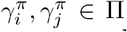 with one being better than the other with respect to all optimization targets. Pareto optimal designs constitute the best tradeoff possible between the studied traits: we can only improve one of them by worsening some other.

We are invested in the minimization of three optimization targets ({*T*_1_, *T*_2_, *T*_3_}):

i. **Error** *ϵ*(*γ*) ∈ [0, 1] in implementing the computational task *φ* (eq. 1). At each evaluation step (*t* = 1, …, *τ*), after damage and regeneration, the network *γ* is fed all 2^*M*^ possible input bits *I_i_* (*i* = 1, …, 2^3^ for *Mux* and *Maj*, figure 1). The network output *O^γ^*(*I_i_*, *t*) is then compared to *φ*(*I_i_*) to compute the average number, over inputs and network lifespan, of mistaken outputs:

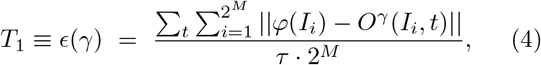

where || · || represents absolute value. We average *T*_1_ over 10 independent realizations of the aging process.
ii. **Connectivity** given by the number of links at *t* = 0 before any damage:

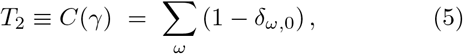

where *δ*_*ω*,0_ is Kronecker’s delta and the sum runs over all possible weights such that each non-zero connection incurs in one unit cost. This target embodies a metabolic burden entailed by wiring costs.
iii. Each network is created with a unique **regeneration rate**:

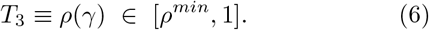

*ρ*(*γ*) specifies the probability with which the network recovers each damaged link at each *t* = 1, …, *τ* of the aging process. A lower bound *ρ^min^* = 0.01 > 0 was chosen as such minimum regeneration power seems to be always present in living systems [16].

The set Γ of possible networks is combinatorially vast. To explore it, we resort to an evolutionary MOO algorithm (figure 2). For a seed (random) population 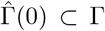 of networks 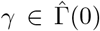 we evaluate their performance in each target {*T*_1_(*γ*), *T*_2_(*γ*), *T*_3_(*γ*)}, select the fittest ones according to a Pareto dominance criterion, and produce diverse offspring through crossover and mutation. This evolves our network ensemble towards the MOO solution Π over generations *g* = 0, …, *g^max^*.

**FIG. 2:**
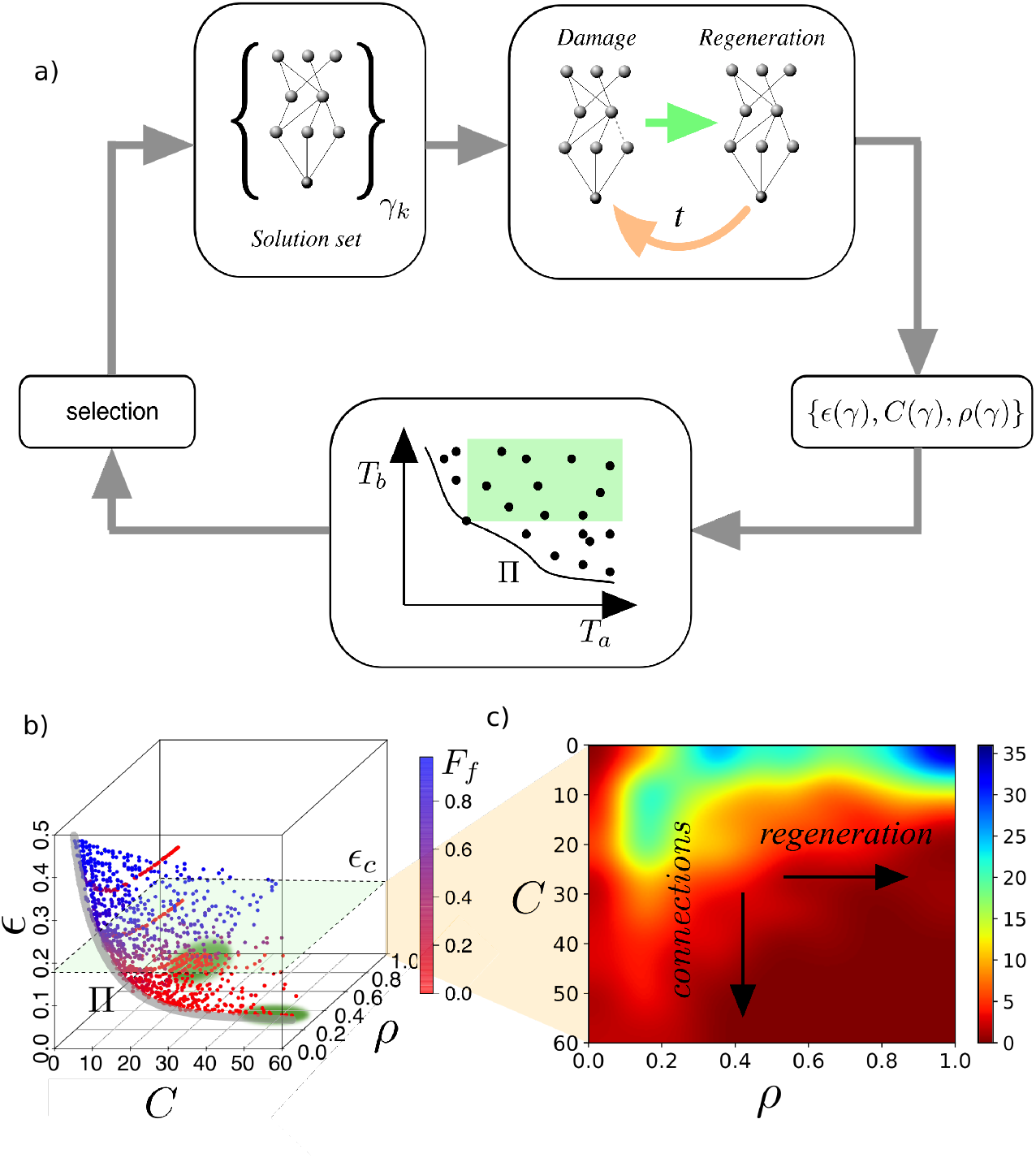
Evolutionary algorithm for neural regeneration and output processing. (a) Iterative process of the MOO algorithm. A population 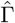 of *N* = 480 networks 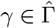 is subjected to a damage: at each of *τ* time steps, their connections are lost with probability *δ*; meanwhile, the network attempts to compute. Computational performance of this damaged network (*T*_1_ ≡ *ϵ*(*γ*)) is measured, along with costs associated to wiring (*T*_2_ ≡ *C*(*γ*)) and regeneration of lost connections (*T*_3_ ≡ *ρ*(*γ*)). These *optimization targets* should be minimized. Bad solutions have a worse performance in all those targets simultaneously (green shaded area): they are *dominated*. Better network designs cannot perform optimally in all senses. They are rather better in trading good performance in a target for worse performance in others, thus avoiding dominance. Less dominated networks at a given iteration of the MOO algorithm are carried to the next generation and used to produce offspring. This algorithm proceeds for *g^max^* iterations. (b) We execute this algorithm ten times for each fixed conditions of damage (*δ*) and length of the aging process (*τ*). This results in a combined population 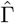 whose non-dominated subset 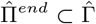 approximates the optimal tradeoff between the targets involved. Blurred green circles indicate extreme phenotypes (investment is maximal in regeneration and minimal in connectivity, or vice versa). A steep relation between the computation error (*T*_1_) and the structural feasibility of the network (as captured by *F_f_*(*γ*), see SM section VII, figures S18-S25) suggests a robust threshold delimiting acceptable computation *ϵ*_*c*_ = 0.2. (c) Density of network designs in 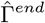 over the *ρ* − *C* plane that compute acceptably (*ϵ*(*γ*) < *ϵ*_*c*_). These plots will be used to explore the tradeoff between regeneration and high connectivity.

Our *target space* (with {*T*_1_, *T*_2_, *T*_3_} as axes, figure 2**b**) allows us to compare networks without favoring performance in any *T_k_* over the others: Given *γ_i_*, *γ_j_* ∈ Γ, we say that *γ_i_ dominates γ_j_* (and note it *γ_i_* ≺ *γ_j_*) if *γ_i_* is no worse than *γ_j_* in all targets (*T_k_*(*γ_i_*) ≤ *T_k_*(*γ_j_*)∀*k* = 1, 2, 3) and it is strictly better in at least one target (∃*k*′, *T*_*k*′_ (*γ_i_*) < *T*_*k*′_(*γ_j_*)).

Using these guidelines (following [43]), we conducted an MOO with a population of *N* = 480 networks over *g^max^* = 4000 generations. At each generation, {*T*_1_(*γ*), *T*_2_(*γ*), *T*_3_(*γ*)} were used to calculate dominance scores: 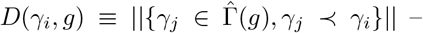 i.e. the number of designs 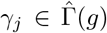 that dominate 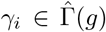. Pareto optimal networks have *D*(*γ^π^* ∈ Π) = 0 (but not the other way around). Poor-performing networks soon become dominated by others. We *copied* all *γ_i_* with *D*(*γ_i_, g*) = 0 into 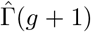. This *elitist* strategy ensures that we never lose the fittest designs. The half of the population with largest *D*(*γ_i_, g*) is discarded. Networks in the remaining half are combined to bring 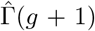 to its full size (*N* = 480). The resulting children are mutated (see Supplementary Material, section III, for further implementation details). We ran this algorithm10 times for each (*δ, τ*) condition. The final population of all 10 runs are combined in a unique set noted 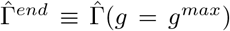. The results reported correspond to the Pareto-optimal networks of this merged data set, 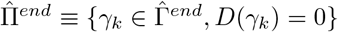 (see Supplementary Material, section IV, to observe all the final results for each (*δ, τ*) condition). We assume 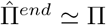, but full convergence cannot be guaranteed.

Figure 2**b** illustrates this for particular (*δ, τ*) conditions. 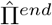 (and actually Π itself) includes designs that compute very badly (large *T*_1_ ≡ *ϵ*(*γ*)) but have been selected because of their negligible regeneration cost and number of links. Pareto dominance offers no principled way to dispose of these networks, even though they would fade away in a biological setting because they plainly fail to function. Interestingly, we observed that computational errors are often associated to deeper structural breakdowns, measured by *F_f_*. Figure 2**b** shows, colorcoded, the feasibility *F_f_*(*γ*) of each network. Those with large computational errors often cannot even convey information from input to output. Plotting *ϵ*(*γ*) vs *F_f_*(*γ*) (see Supplementary Material, section VII) we noted that this degradation of computation capabilities and network structure follows a logistic curve with a marked threshold *ϵ_c_* ~ 0.24. This value turned out to be similar across realizations of the algorithm and for different (*δ, τ*) conditions (both for *Mux* and *Maj* functions; see Supplementary Material, figures S18 to S25). We took it as a heuristic limit (horizontal plane in figure 2**b**) to select *acceptably working* designs (we took *ϵ_c_* = 0.2 to be on the safe side). Figure 2**d** shows all networks with a performance better than this threshold (*T*_1_ ≡ *ϵ*(*γ*) < *ϵ*_*c*_ = 0.2) in a *C* − *ρ* map. (See Supplementary Material, section V, to observe the density plots for each (*δ, τ*) condition, section I, to check on all the parameters of the model, and section III to check on further implementation details.)

## III. RESULTS

### A. Tradeoff between connectivity and regeneration

Figure 3 shows abundance of networks 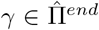 with *ϵ*(*γ*) < *ϵ_c_* ~ 0.2 (i.e. Pareto optimal designs that further satisfy the reasonable phenotypic performance *ϵ_c_*, which also entails a persisting structural integrity – large feasibility *F_f_* (*γ*)) for a low damage *δ* condition, both for *Mux* and *Maj* as computational tasks. The plotted abundance of designs across the *ρ* − *C* plane captures the trade-off between connectivity (*T*_2_ ≡ *C*(*γ*)) and regeneration (*T*_3_ ≡ *ρ*(*γ*)). A broad range of optimal solutions compatible with proper functionality is showcased.

**FIG. 3:**
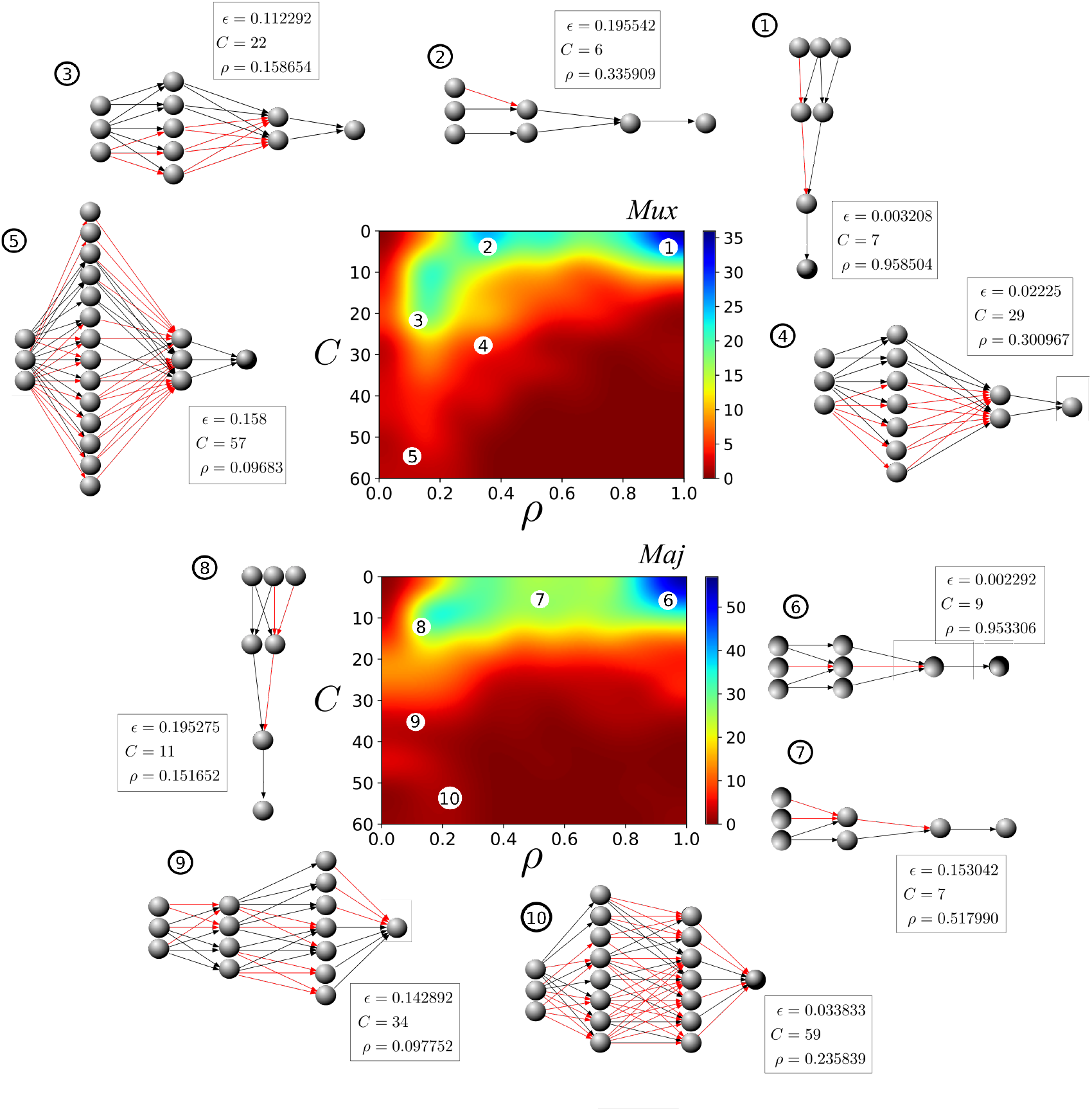
Connectivity-regeneration (*C* − *ρ*) tradeoff as a means to achieve reliable computation. Two selected density plots for *Mux* (top) and *Maj* (bottom) of solutions 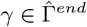 with reliable computation (*ϵ*(*γ*) < *ϵ*_*c*_) throughout the *ρ* − *C* plane. Alongside, representative networks from characteristic regions of phenotype space. Performance of these networks in all three targets is shown (enclosed squares). Results obtained for *δ* = 0.04, *τ* = 300 for *Mux* and *δ* = 0.02 and *τ* = 1500 for *Maj*. (See SM for other values.)

Both panels (and further plots in SM) show a higher density of solutions around areas with less connections and higher regeneration (e.g. peak in the upper right corner, figure 3**a**). Graphs labeled 1 and 2 for *Mux* (figure 3**a**) and 6, 7, and 8 for *Maj* (figure 3**b**) illustrate minimal circuits implementing those tasks. The density of designs found with lower regeneration rates (which demands higher connectivity) is notably smaller in comparison. Some graphs (3, 4, and 5 for *Mux*; 9 and 10 for *Maj*) sample this more sparsely occupied region with lower regeneration levels and more densely connected circuits. Trends are shared among the two Boolean functions tested, such as the lower density of solutions in this region of phenotype space, and the resilience that is obtained trough abundant, duplicated links. This hints us at general patterns emerging despite the potential variety in phenotypes imposed by different computational tasks.

### B. Network lifespan and external damage act as evolutionary pressures

As discussed above, our aim is to asses the influence of additional distinct features on our evolutionary framework, namely the lifespan of networks and the damage inflicted to their links during the aging process. The former offers a window to the effects of life length (*τ*) on the selection process whereas the latter, implemented by *δ*, allows including the stochastic perturbations that hamper proper computation. What is the impact of lifetime (*τ*) or damage rates (*δ*) in the optimal space of solutions resulting from our model?

We have found that, for a fixed damage rate, longer agent life times act as an evolutionary pressure, such that the algorithm better explores the optimal tradeoff by pushing harder our evolving population against it. The same happens in the opposite case, e.g. increasing damage rates for fixed life times. To capture this, we compute the amount of non-dominated versus dominated solutions in 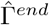:

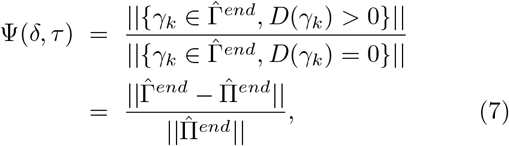

where *D*(·) is again the dominance score. Each of our evolution takes place under fixed (*δ, τ*) conditions. Those instances that result in larger Ψ retain more network designs that are sub-optimal (in the Pareto sense) with respect to other surviving designs. This is, such (*δ, τ*) settings are less strict during selection, such that networks that perform relatively worse in all targets simultaneously are still retained. Opposed to this, (*δ, τ*) values resulting in a smaller Ψ are more severe regarding Pareto optimality selection – i.e. evolutionary pressure to select Pareto optimal solutions is higher. See Fig. 4a to check the effect of fixed damage rates and increasing life times, and Fig. 4b to illustrate the effect of fixed lifetime and increasing damage. However, both effects are present in both plots. (Check Supplementary Material, section VI, to check the rest of parameter values either for *Mux* and *Maj*.)

**FIG. 4:**
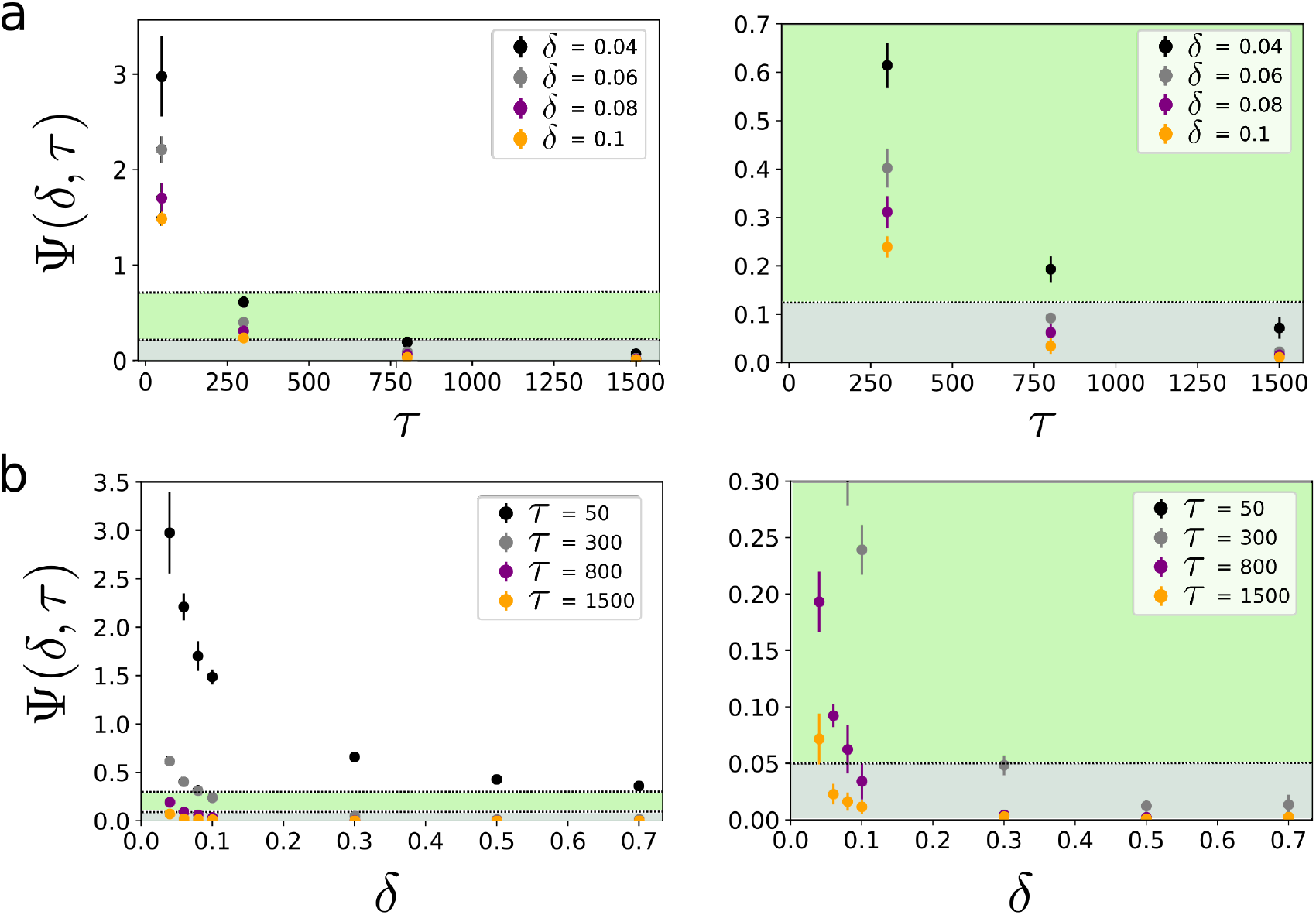
Life time *τ* and damage rate *δ* act as evolutionary pressures. (a) Ratio Ψ(*δ, τ*) between the number of non-Pareto and Pareto-optimal solutions in 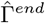 as a function of the *τ* for fixed values of damage rate *δ* (for *Mux*). Ψ decreases for increasing *τ*. Only a sample of *δ* values (0.04, 0.06, 0.08 and 0.1) is shown to help visualization (other conditions, in SM section VI, also for *Maj*; all results in Supplementary Material support the conclusions in the main text). (b) Same ratio for fixed values of *τ* and varying *δ*. Again, Ψ drops as *δ* increases. Decreases in Ψ capture a higher evolutionary pressure towards Pareto optimality.

Which are the reasons for such observation? Despite both network lifespan and damage rate exert a pressure of the same nature (as measured by Ψ(*δ, τ*), the ultimate reasons for such observation might be different. An explanation for large *τ* as an evolutionary pressure for Pareto optimality could lay in the fact that longer life times are synonym of dealing with more extended and numerous threats entailing a larger information retrieval from the environment. Therefore, improvements during evolutionary time can have a larger impact on this longer living populations compared to shorter living ones. Regarding the influence of the degree of damage inflicted to networks (*δ*), the reason for such evolutionary pressure is probably rooted to the inherent harsher survival conditions imposed in larger damage regimes.

Importantly, the interplay of both features might also be acting as an evolutionary pressure (as measured by Ψ(*δ, τ*)). As presented in section IIC, damage rates and network lifespans can be collapsed into a dimensionless ratio, *r_τ_* ≡ *δ* · *τ* ~ *τ/τ*_*link*_, as the lifespan of the connections *τ_link_* grows monotonously with 1/*δ*, thus capturing a relationship between time-scales proper of whole organisms versus those of its parts. An increase of this ratio also entails lower values of Ψ(*δ, τ*), meaning that an increase in the difference between the lifespan of the components versus that of the organism might be playing a role in the described observations.

### C. Damage rates influence the overall shape of the optimal tradeoff

The resulting 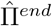 embody the optimal tradeoff between all targets involved. This optimal tradeoff can be visualized if plotted in target space, where its shape can be different for each (*δ, τ*) fixed conditions. Our exhaustive exploration of (*δ, τ*) combinations (see SM section IV) shows a clear overall change in the shape of this optimal tradeoff as damage increases – a change, furthermore, that has consequences in terms of phenotype space accessibility and exploration as we discuss below. Figure 5**a-c** shows networks in 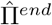 for increasing damage regimes. Designs with *ϵ*(*γ*) > *ϵ*_*c*_ are retained here. 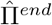 appears smooth for the low damage regime (*δ* = 0.01), meaning that the optimal tradeoff does not present large cavities or singular points when plotted in target space (figure 5**a**). This surface becomes more rugged (i.e. its curvature changes, thus generating cavities) for increasing damage rates (*δ* = 0.1, 0.7, figure 5**b, c**).

**FIG. 5:**
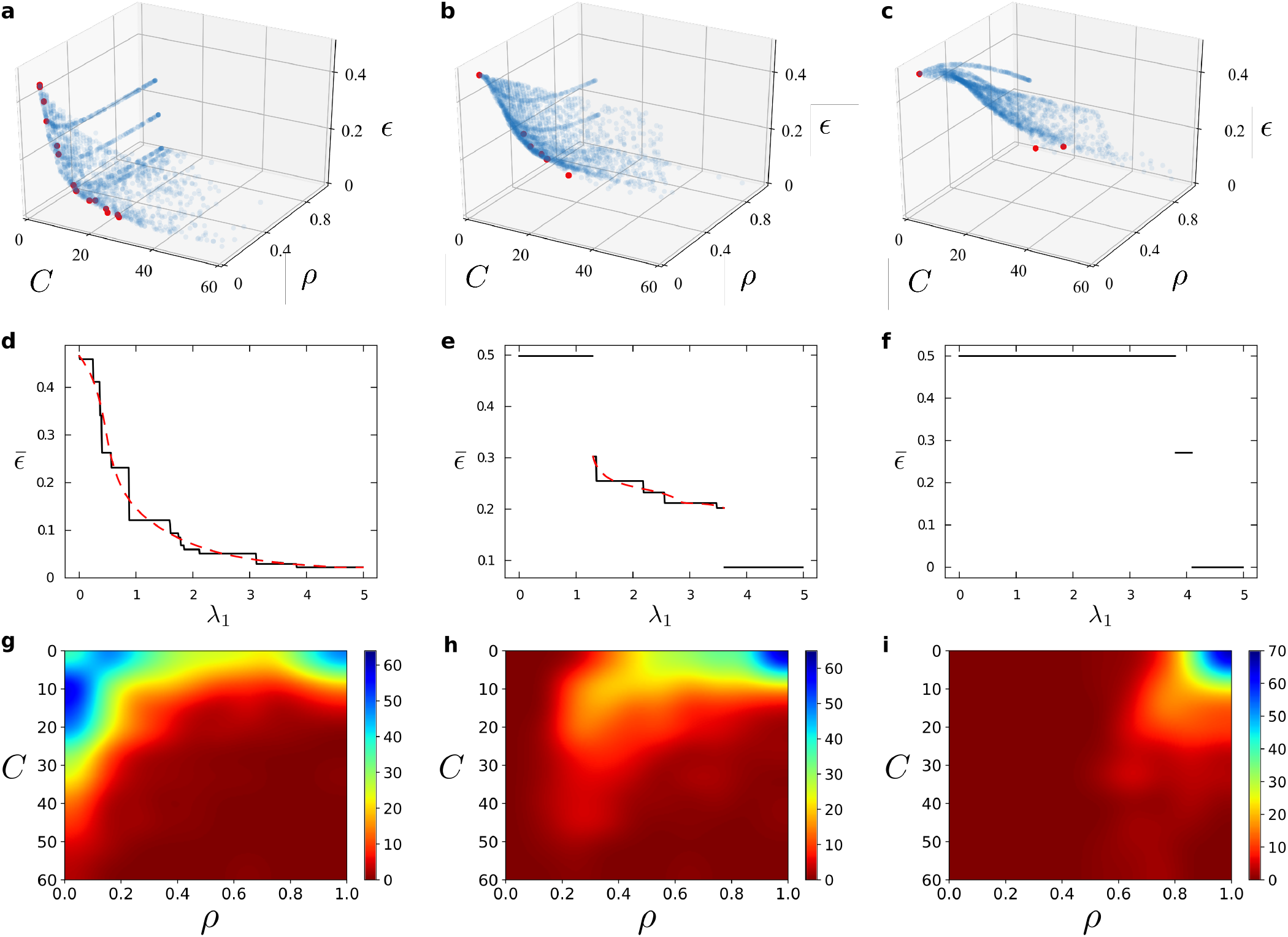
Overall shape of the optimal tradeoff and accessibility of phenotype space. Pareto optimal strategies 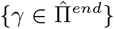 under fixed conditions ((a) *δ* = 0.01, *τ* = 1500; (b) *δ* = 0.1, *τ* = 300; (c) *δ* = 0.7, *τ* = 50) plotted in *target space*. Each blue dot represents {*T*_1_(*γ*), *T*_2_(*γ*), *T*_3_(*γ*)} for a given 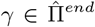. The shape of each embedding surface determines how accessible phenotype space is if minimizers of a global utility function Ω = Σ λ_*i*_*T*_*i*_ were sought. Red dots represent such global optima as *λ*_2_ = 0.009 and *λ*_3_ = 2 are kept fixed and *λ*_1_ ∈ [0, 5]. Increasing damage ((a) *δ* = 0.01, (b) *δ* = 0.1, (c) *δ* = 0.7) results in increasingly more rugged tradeoffs. Smooth tradeoffs (a) are sampled more evenly by global optimizers, so that successive global optima are similar to each other. Whatever property we plot of them, e.g. their computational error *ϵ*(*γ*) (d, solid black lines), varies relatively smoothly with *λ*_1_. Some discreteness remains due to the numerical nature of our experiments (dashed red lines illustrate the underlying continuous dependency). The corresponding density plot (g) with solutions fulfilling *ϵ*(*γ*) < *ϵ*_*c*_ shows the tradeoff *ρ* − *C* available for the low damage regime (already shown in figure 3). Tradeoffs with cavities (b, c) leave some phenotypes unsampled by global optima. These often take leaps in phenotype space, resulting in sudden jumps in any measured property of such solutions as a function of *λ*_1_ (e, f). These plots show how a heightened damage rate has an ability to hamper access to existing phenotypes and fracture the continuity of global optimal designs. The corresponding density plots (h, i) with solutions fulfilling *ϵ*(*γ*) < *ϵ*_*c*_ show the constraints imposed by increasing damage regimes in the *C* − *ρ* projection. Crucially, these available designs can be further constrained by the fracture of the continuity of global optimal designs. See S2-S7 in SM for further evidence linking ruggedness of the optimal tradeoff to increased damage rate, both for *Mux* and *Maj* functions.

This varying ruggedness of optimal tradeoffs can tell us something about how accessible our space of optimal solutions is. We could weight all targets linearly into a global optimization function: Ω(*γ*, Λ) = *λ*_1_*T*_1_(*γ*) + *λ*_2_*T*_2_(*γ*) + *λ*_3_*T*_3_(*γ*), where Λ ≡ {*λ*_1_, *λ*_2_, *λ*_3_} represent explicit evolutionary biases towards a specific target. For example, a large *λ*_2_ versus low *λ*_1,3_ indicates that, in a given environment, minimizing the initial metabolic cost of links is outstandingly important. Giving values to the *λ_k_* could thus define a specific set of external environmental pressures. Then the minimization of Ω(*γ*, Λ) selects one single solution 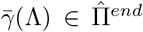 out of the Pareto optimal tradeoff. This is, the imposition of such specific constraints would force evolution towards a determined region of phenotype space.

Cavities and singular points in 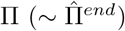 have been linked to phase transitions [37–41] and critical phenomena [39, 41] that arise as the *λ_k_* are varied – i.e. as potential biases in a niche change (perhaps over time, or perhaps because the species has left that niche). A phase transition in our problem would indicate that certain network designs are persistently stable under a range of external evolutionary conditions, and that a switch from a network design to another would sometimes be drastic even if those external conditions would vary just slightly.

Such cavities and singular points are absent for the low *δ* example in figure 5**a** (the same regime was shown in figure 3), meaning that the whole phenotypic space of nervous system maintenance strategies can be smoothly visited as evolutionary pressures vary. To illustrate this we have set *λ*_2_ and *λ*_3_ to a fixed value while varying *λ*_1_ ∈ [0, 5]. This means that we sample evolutionary pressures that initially disregard good computation but progress towards situations in which computing correctly becomes more pressing. Network designs 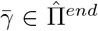 that minimize Ω(*γ*, Λ) for the range of *λ*_1_ are displayed in red in figure 5**a**. Figure 5**d** shows the computational error for these absolute optima 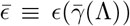 as a function of *λ*_1_. The numerical nature of our experiments introduces some unavoidable discreteness in 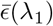); but overall we can see how 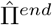 is rather smoothly sampled (as compared to the next cases) and optimal solutions progress parsimoniously into each other as external evolutionary pressures change.

Additionally, figure 5**a** shows how 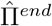 becomes virtually flat for *T*_1_ ≡ *ϵ* → 0. Decrementally small improvements in computation can only be achieved by increasingly larger investment on *T*_2_ ≡ *C* and *T*_3_ ≡ *ρ*. Perfect computation (*ϵ* = 0) could be reached ultimately. But this plot reveals a decreasing return for low errors. Organisms eventually need exaggerated investments to achieve negligible computational improvements. This is reflected by a sparse sampling of those costly areas of phenotype space, also reflected in the plots of abundance of designs across the *ρ* − *C* plane (figure 3).

Optimal tradeoffs present cavities for the examples in larger damage regimes (figures 5**b, c**). Red dots show absolute optima as the same values of Λ are sampled. The cavities in 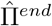 entail that a lower number of different absolute optima are recovered for *δ* = 0.1, and even less for *δ* = 0.7 under the same procedure. This is so because certain phenotypes remain optimal over a wide range of *λ*_1_, thus preventing us from visiting part of phenotypic space. Furthermore, when such paramount designs stop being global optima, the new preferred network is found far apart in 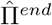. A small change in *λ*_1_ would then demand a prompt yet drastic adaptation to accommodate the new best. This is reflected in 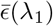 plots (figures 5**e-f**) through huge improvements in performance as the bias towards minimizing the computational error (*λ*_1_) increases.

Notice that in these cases (figures 5**b, c**) the overall shape of the Pareto optimal front is also constraining to a much reduced area the set of designs which retain reliable computations (*ϵ* < *ϵ*_*c*_, see the net effect on the *ρ* − *C* density plots in figures 5**h, i**), if compared with the low damage rate regime shown in figures 3 and 5**a,g**. Thus, increased damage (besides constraining the available phenotypic space due to emerging phase transitions) also results in a much more narrower space when looking only at *acceptably performing* designs, disregarding of any global optimization of Ω.

Importantly, it cannot be discarded that either the network lifespan (*τ*) or the interplay between it and the lifespan of the connections (*τ_link_*), measured through *r_τ_*, may also have an influence on the change in the overall shape of optimal tradeoffs (*r_τ_* = 15, 30, 35 in figures 5**a,b,c**, respectively). However, the observations retrieved from the set of tested parameters (see Supplementary Material, section IV) show that the clear driver of the observed effect is the increase in damage rates regimes.

## IV. DISCUSSION

In this paper we attempt to provide insights on which are the evolutionary pressures or drivers that may underpin the evolution of nervous systems and its regeneration capabilities. Due to the generality of the model, we do not aim to answer specific questions but rather extract general principles which might pave the way to future research directions. The simplifications we make are considerable, starting from the use of artificial feed-forward neural networks to represent a nervous system. Regarding the dynamics of the system, the so-called aging process to which our networks are exposed and their subsequent response is a simplification of the processes of axonogenesis and neurogenesis [16] observed in real nervous systems. While only axonogenesis is explicitly incorporated in the model, neurogenesis can be considered to be present as a secondary response linked to the recovery of connections (whose loss can cause the breakdown of neuron functionality). On the other hand, it is also known that regeneration response is dependent on anatomical location and type of damage [16], but such a feature has not been incorporated in any way, being another simplification of the model. And yet our schematic framework retains what we think are some key factors to understand the evolutionary drivers of neural systems maintenance. We can thus disentangle some pressures at play as well as isolate relevant factors for future studies. This would probably be more difficult with more complicated models.

First, our results show that, in the domain of reliable computation, a tradeoff might be at play between high density of connections and high regeneration capabilities. This tradeoff shapes the phenotypic space of possible designs for network maintenance, and it reminds us of the variety of such strategies in the natural world. The region of phenotype space with higher connectivity presumably achieves robust computation through specific redundant connections or degenerate mechanisms [44, 45]. The alternative, which relies on large regeneration rates, is found in the region of phenotype space where networks are smaller. We observe that these are also the networks preferentially explored by our MOO algorithm. This is so even if, intuitively, a higher number of connections could result in a potentially larger amount of different architectures that solve a same task.

This suggests that regeneration can be a more reliable strategy – at least at the scales explored. Strategies betting on duplicate pathways might need to deal, e.g., with emerging interactions between surviving connections as faulty ones are removed. Such problems could be bypassed by alternate computational paradigms – e.g. distributed computation [46, 47] or reservoir computing [48–52]. The latter can take explicit advantage of such emergent behaviors, so that damage and regeneration properties might be an evolutionary bias towards biological solutions based on reservoir computing [53]. Such alternatives could unlock further regions of phenotypic space with a high number of similar solutions. But such computational strategies depend crucially on huge numbers of units, and might be unreachable for our experiments; thus returning us to the observed bias towards regeneration.

Secondly, we have found that both the extent of external damage and the network lifespan are acting as active evolutionary pressures over the whole population so that it eventually contains more Pareto optimal individuals. The increase of both factors gives rise to more strict evolutionary scenarios, in which the pressure to select Pareto optimal solutions is higher, resulting in more diverse optimal strategies (better exploration of the optimal tradeoffs). Interestingly, the relationship *r_τ_* between organismal lifetimes (*τ*) and the timescale of its component parts (*τ_link_* ~ 1/*δ*) is also suggested to be acting as an evolutionary pressure towards this direction. Focus on this ratio becomes a very enticing research avenue if we wonder at what level (organismal versus component part) can a Darwinian process store the information gathered as evolution proceeds.

Third, from the observations retrieved we have found that regimes of increasing damage (which can be considered an ecological factor) result in more rugged tradeoffs. Such tradeoffs present cavities, which is the characteristic signature of first order phase transitions [37–41]. Under varying evolutionary biases, the presence of such transitions can result in history dependency effects similar to hysteresis [54]. In such phenomena, evolutionary pressures confronted by a species would vary just slightly and yet a preferred global optimum would change drastically. The species might need to adapt swiftly and retain suboptimal aspects due to frozen accidents, thus resulting in increased evolutionary path dependency.

All of this suggests that i) discrete phenotypic space, drastic changes expected under varying external evolutionary pressures, iii) phenotypic space becoming less accessible, and iv) heightened path dependency in evolution should all become more prominent as the external erosion of our system is higher. Overall, this would mean that higher damage rates could induce organisms to commit to specific nervous systems maintenance strategies, potentially renouncing an ability to switch options with relative ease. Damage could change swiftly as living beings migrate to more benign environments or are suddenly locked on harsher conditions. Similarly to an inherent shorter life-time of component parts, a heightened external damage rate has the ability to harshen the strictness of the selection process (again, in a Pareto optimality sense; thus implying population-wide phenomena) and also of promptly shifting the shape of the optimal tradeoff. Future work will be required to explore these results in a more general context of multicellularity (natural and synthetic) where cognitive complexity is a well-defined dimension [55].

## Acknowledgments

The authors thank the members of the Complex Systems Lab for useful discussions, specially Jordi Piñero. This work has been supported by an ERC Advanced Grant Number 294294 from the EU seventh framework program (SYNCOM), by the Botín Foundation by Banco Santander through its Santander Universities Global Division, a MINECO grant FIS2015-67616 fellowship cofunded by FEDER/UE, by the Universities and Research Secretariat of the Ministry of Business and Knowledge of the Generalitat de Catalunya and the European Social Fund, and by the Santa Fe Institute. L.F.S. has also been supported by the IFISC (Institute of Interdisciplinary Physics and Complex Systems) as part of the María de Maeztu Program for Units of Excellence in R&D (Grant No. MDM-2017-0711).

## Notes

#### Summary of Updates

Minor changes in text, minor corrections in figures, funding institution added.

## References

[1] Jablonka E, Lamb MJ. 2006 The evolution of information in the major transitions. J. Theor. Biol. 239(2), 236–246.

[2] Bickerton D. 1990 Language and species, Ch. 4. University of Chicago Press.

[3] Von Neumann J. 1956 Probabilistic logics and the synthesis of reliable organisms from unreliable components. Automata Studies 34, 43–98.

[4] Winograd S, Cowan JD. 1963 Reliable computation in the presence of noise. MIT Press, Cambridge, Mass.

[5] Barlow HB, Levick WR. 1965 The mechanism of directionally selective units in rabbit’s retina. J. Phys. 178(3), 477–504.

[6] Gavrilov LA, Gavrilova NS. 2001 The reliability theory of aging and longevity. J. Theor. Biol. 213(4), 527–545.

[7] Ferretti P. 2011 Is there a relationship between adult neurogenesis and neuron generation following injury across evolution? Eur. J. Neurosci. 34(6), 951–962.

[8] Moffet SB. 2012 Nervous system regeneration in the invertebrates. Springer Scoence & Business Media.

[9] Levin M. 2007 Large-scale biophysics: ion flows and regeneration. Trends Cell Biol. 17(6), 261–270.

[10] Levin M. 2009 Bioelectric mechanisms in regeneration: unique aspects and future perspectives. In Seminars in cell & developmental biology 20(5), 543–556. Elsevier.

[11] Tseng A-S, Beane WS, Lemire JM, Masi A, Levin M. 2010 Induction of vertebrate regeneration by a transient sodium current. J. Neurosci. 30(39), 13192–13200.

[12] Eriksson PS, Perfilieva E, Björk-Eriksson T, Alborn AM, Nordborg C, Peterson D, Gage FH. 1998 Neurogenesis in the adult human hippocampus. Nat. Med. 4(11), 1313.

[13] van Praag H, Schinder AF, Christie BR, Toni N, Palmer TD, Gage FH. 2002 Functional neurogenesis in the adult hippocampus. Nature 415(6874), 1030.

[14] Hack MA, Saghatelyan A, Chevigny A, Pfeifer A, Ashery-Padan R, Lledo P-M, Gotz M. 2005 Neuronal fate determinants of adult olfactory bulb neurogenesis. Nature Neurosci. 8(7), 865.

[15] Zupanc GKH. 2006 Neurogenesis and neuronal regeneration in the adult fish brain. J. Comp. Physiol. A 192(6), 649.

[16] Tanaka EM, Ferretti P. 2009 Considering the evolution of regeneration in the central nervous system. Nat. Rev. Neurosci. 10(10), 713.

[17] Chen BL, Hall DH, Chklovskii DB. 2006 Wiring optimization can relate neuronal structure and function. Proc. Nat. Acad. Sci. 103(12), 4723–4728.

[18] Chklovskii DB, Schikorski T, Stevens CF. 2002 Wiring optimization in cortical circuits. Neuron 34(3), 341–347.

[19] Raj A, Chen Y-H. PLoS one 6(9), e14832.

[20] Miller GF, Todd PM, Hegde SU. 1989 Designing Neural Networks using Genetic Algorithms. In ICGA 89, 379–384.

[21] Angeline PJ, Saunders GM, Pollack JB. 1994 An evolutionary algorithm that constructs recurrent neural networks. IEEE T. Neural Net. 5(1), 54–65.

[22] Yao X. 1999 Evolving artificial neural networks. Proc. IEEE 87(9), 1423–1447.

[23] Floreano D, Dürr P, Mattiussi C. Neuroevolution: from architectures to learning. Evolutionary Intelligence, 1(1), 47–62.

[24] Stanley KO, Miikkulainen R. 2002 Evolving neural networks through augmenting topologies. Evol. Comput. 10(2), 99–127.

[25] Stanley KO, D’Ambrosio DB, Gauci J. 2009 Artif. Life 15(2), 185–212.

[26] Eckmann J-P, Feinerman O, Gruendlinger L, Moses E, Soriano J, Tlusty T. 2007 The physics of living neural networks. Phys. Rep. 449(1-3), 54–76.

[27] Székely G. 1965 Logical network for controlling limb movements in urodela. Acta Physiol. Hung. 27, 285.

[28] Stent GS, Kristan WB, Friesen WO, Otto CA, Poon M, Calabrese RL. 1978 Neuronal generation of the leech swimming movement. Science 200(4348), 1348–1357.

[29] Friesen WO, Stent GS. 1978 Neural circuits for generating rhythmic movements. Annu. Rev. Biophys. Bio. 7(1), 37–61.

[30] Amari S-I. 1977 Dynamics of pattern formation in lateral-inhibition type neural fields. Biol. Cybern. 27(2), 77–87.

[31] Chen J, Mouret J-B, Lipson H. 2013 The evolutionary origins of modularity. Proc. R. Soc. B 280(1755), 20122863.

[32] Floreano D, Nolfi S, Mondada F. 2001 Co-evolution and ontogenetic change in competing robots. Advances in the evolutionary synthesis of intelligent agents, 273–306.

[33] Pfeifer R, Lungarella M, Iida F. 2007 Self-organization, embodiment, and biologically inspired robotics. American Association for the Advancement of Science 318(5853), 1088–1093.

[34] Coello C, Zacatenco C. 2006 Twenty years of evolutionary multi-objective optimization: A historical view of the field. IEEE Comput. Inel. M. 1(1), 28–36.

[35] Schuster P. 2012 Optimization of multiple criteria: Pareto efficiency and fast heuristics should be more popular than they are. Complexity 18(2), 5–7.

[36] Seoane LF. 2016 Multiobjetive optimization in models of synthetic and natural living systems. Doctoral dissertation, Universitat Pompeu Fabra.

[37] Seoane LF, Solé R. 2013 A multiobjective optimization approach to statistical mechanics. arXiv preprint arXiv:1310.6372.

[38] Seoane LF, Solé R. 2015 Phase transitions in Pareto optimal complex networks. Phys. Rev. E 92(3), 032807.

[39] Seoane LF, Solé R. 2015 Systems poised to criticality through Pareto selective forces. arXiv preprint arXiv:1510.08697.

[40] Seoane LF, Solé R. 2016 Multiobjective optimization and phase transitions. In Proceedings of ECCS 2014. 259–270.

[41] Seoane LF, Solé R. 2018 The morphospace of language networks. Sci. Rep. 8.

[42] Rojas R. 2013 Neural networks: a systematic introduction. Springer Science & Business Media.

[43] Konak A, Coit DW, Smith AE. 2006 Multi-objective optimization using genetic algorithms: A tutorial. Reliab. Eng. Syst. Safe. 91(9), 992–1007.

[44] Tononi G, Sporns O, Edelman GM. 1999 Measures of degeneracy and redundancy in biological networks. Proc. Nat. Acad. Sci. 96(6), 3257–3262.

[45] Edelman GM, Gally JA. 2001 Degeneracy and complexity in biological systems. Proc. Nat. Acad. Sci. 98(24), 13763–13768.

[46] Macía J, Posas F, Solé R. 2012 Distributed computation: the new wave of synthetic biology devices. Trends Biotechnol. 30(6), 342–349.

[47] Regot S, Macía J, Conde N, Furukawa K, Kjellén J, Peeters T, Hohmann S, De Nadal E, Posas F, Solé R. 2011 Distributed biological computation with multicellular engineered networks. Nature 469(7329), 207.

[48] Jaeger H. 2001 The “echo state” approach to analysing and training recurrent neural networks-with an erratum note. In Bonn, Germany: German National Research Center for Information Technology GMD Technical Report 148(34), 13.

[49] Maass W, Natschläger T, Markram H. 2002 Real-time computing without stable states: A new framework for neural computation based on perturbations. Neural Comput. 14(11), 2531–2560.

[50] Jaeger H, Maass W, Principe J. 2007 Special issue on echo state networks and liquid state machines. Elsevier Science.

[51] Verstraeten D, Schrauwen B, d’Haene M, Stroobandt D. 2007 An experimental unification of reservoir computing methods. Neural Networks 20(3), 391–403.

[52] Lukoševičius M, Jaeger H, Schrauwen B. 2012 Reservoir computing trends. KI-Künstliche Intelligenz 26(4), 365–371.

[53] Seoane LF. 2019 Evolutionary aspects of reservoir computing. Philos. T. R. Soc. B 374(1774), 20180377.

[54] Solé R. 2011 Phase Transitions. Princeton U. Press.

[55] Ollé-Vila A, Duran-Nebreda S, Conde-Pueyo N, Montáñez R, Solé R. 2016 A morphospace for synthetic organs and organoids: the possible and the actual. Integr. Biol. 8(4), 485–503.

